# Mapping-free variant calling using haplotype reconstruction from k-mer frequencies

**DOI:** 10.1101/153619

**Authors:** Peter Audano, Shashidhar Ravishankar, Fredrik Vannberg

## Abstract

1

**Motivation:** The standard protocol for detecting variation in DNA is to map millions of short sequence reads to a known reference and find loci that differ. While this approach works well, it cannot be applied where the sample contains dense variants or is too distant from known references. *De novo* assembly or hybrid methods can recover genomic variation, but the cost of computation is often much higher. We developed a novel k-mer algorithm and software implementation, Kestrel, capable of characterizing densely-packed SNPs and large indels without mapping, assembly, or de Bruijn graphs.

**Results:** When applied to mosaic penicillin binding protein (PBP) genes in *Streptococcus pneumoniae,* we found near perfect concordance with assembled contigs at a fraction of the CPU time. Multilocus sequence typing (MLST) with this approach was able to bypass *de novo* assemblies. Kestrel has a very low false-positive rate when calling variants over the whole genome, but limitations of a purely k-mer based approach affect sensitivity.

**Availability:** Source code and documentation for a Java implementation of Kestrel can be found at https://github.com/paudano/kestrel. All test code for this publication is located at https://github.com/paudano/kescases.

## 2 Introduction

Although modern alignment tools are designed to handle errors and variation, a sequence read that differs significantly from the reference cannot be confidently assigned to the correct location. When these reads are mapped, the low alignment confidence leads to low variant call confidence, and it becomes difficult to separate true variants from false calls. In some regions, the read may be clipped or not mapped at all. As a result, variant calling algorithms that rely solely on alignments cannot characterize these events.

There are several alternative approaches for poor alignments. Calling variants from *de novo* assembled contigs may work for monoploid organisms, but since it reduces the read coverage to 1, false-calls cannot easily be corrected (Olson *et al.,* 2015). Building the de Bruijn graphs that are typically employed in assembly may also require many gigabytes of memory or computing clusters (Li *et al.,* 2010). Cortex (Iqbal *et al.,* 2012) was designed to overcome limitations of the *de novo* assembly approach, but it still relies de Brujin graphs. Platypus (Rimmer *et al.,* 2014) performs local assemblies, however, it incurs the overhead of both read mapping and local assembly if variant loci are not known *a priori.*

One possible approach is to use the information contained within a set of k-mer frequencies over the sequence data. K-mers can be represented numerically, they do not rely on sequence alignments, and k-mer counting methods are fast. To date, sparsely-spaced SNPs have been corrected with such an approach (Gardner and Slezak, 2010; Gardner and Hall, 2013), but a robust variant calling algorithm has never been shown to work. Because dense SNPs and large insertions can create many k-mers very different from any reference sequence, how such an approach could work efficiently is not immediately clear.

We constructed an algorithm that relies strictly on k-mer frequencies over sample sequence data and ordered k-mers in a reference sequence to call variants. This method first finds regions of variation over the reference using disruptions in the frequency distribution of expected k-mers, and by beginning with unaltered k-mers at the flanks of such regions, it can greatly simplify the search through k-mer space while resolving variation.

## 3 Methods

### 3.1 Algorithm overview

Using features in KAnalyze (Audano and Vannberg, 2014) version 2.0.0, the frequency of each k-mer in the sample is stored in an indexed k-mer count file (IKC) (**Fig. 1(a)**). The IKC records are grouped by a k-mer minimizer (Roberts *et al.,* 2004) and sorted. This structure tends to cluster k-mers from the original sequence, and by reading it as a memory mapped file, k-mers can be rapidly queried with low memory requirements using search optimizations similar to those implemented in Kraken (Wood and Salzberg, 2014).

**Figure 1:**
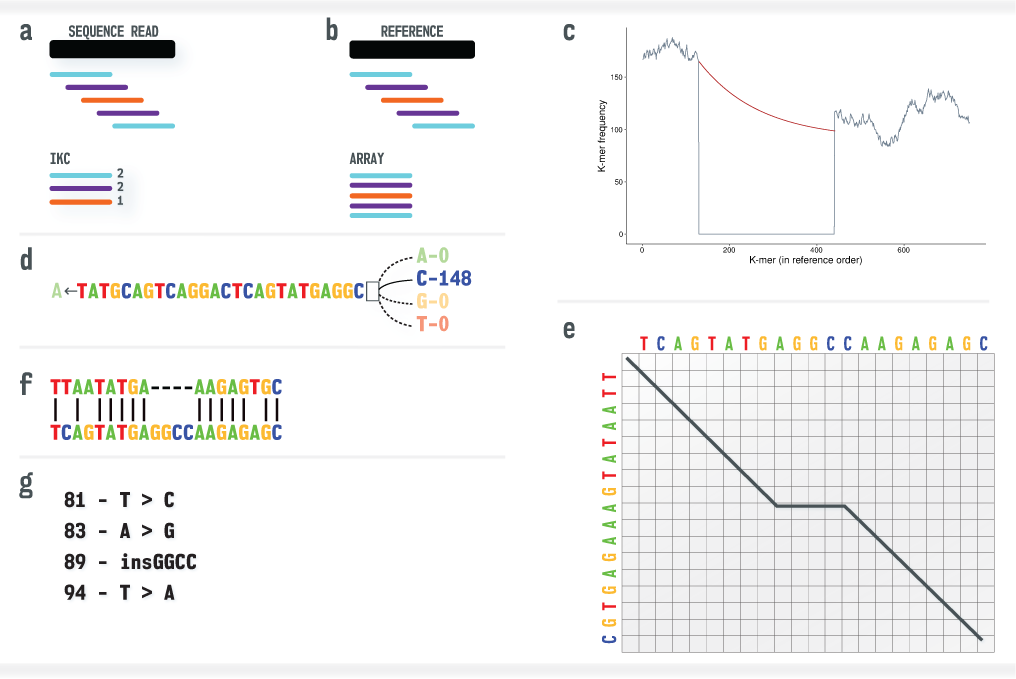
Overview of the Kestrel process from sequence data to variant call. (**a**) The sequence reads are converted to an IKC file. (**b**) The reference sequence is converted into an array of k-mers and left in reference order. (**c**) K-mer frequencies from the sequence reads (vertical axis) are assigned to the ordered k-mers of the reference (horizontal axis). A decline and recovery of the frequencies bound an active region where one or more variants are present. The recovery threshold is degraded with an exponential decay function (red) to allow for declining read coverage. (**d**) Starting from the left anchor k-mer (last k-mer with a high frequency), the first base is removed, each possible base is appended, and the base that recovers the k-mer frequency is appended to the haplotype. (**e**) A modified alignment algorithm tracks haplotype reconstruction and terminates the process when an optimal alignment is reached. (**f**) This algorithm yields an alignment of the reference sequence and haplotype within the active region. (**g**) Variant calls are extracted from the alignment.

From the reference sequence, the frequencies for each k-mer and its reverse complement are obtained from the IKC file and summed, but these are left in order (**Fig. 1(b)**). Kestrel searches the resulting array for loci where the frequency declines and recovers (**Fig. 1(c)**), which suggests the k-mers of the reference and sample differ. Analogous to the GATK (McKenna *et al.,* 2010) HaplotypeCaller, this low-frequency region is called an *active region,* and *haplotypes* are reconstructed over it.

Starting with the high-frequency k-mer immediately to the left of the active region, the first base is removed, all four bases are appended to this (k - 1)-mer, and the k-mer frequency is queried for each of the four possibilities (**Fig. 1(d)**). An equivalent process is performed on the reverse complement k-mer, and the frequencies are summed. The base that yields a high frequency k-mer is appended to the haplotype. The new k-mer ending in that base is then used to find the next base by the same process. If more than one of these k-mers has a high frequency, a haplotype is assembled for each one.

A modified Smith-Waterman (Smith and Waterman, 1981) algorithm guides the process by aligning the active region and the haplotype as it is reconstructed (**Fig. 1(e)**). By setting an initial score and disallowing links to zero score states, alignments are anchored on the left and are allowed to extend until an optimal alignment is obtained. Variant calls follow trivially from the alignment (**Fig. 1(f)** and **Fig. 1(g)**). By relaxing these criteria and by building in either the forward or reverse direction, it is possible to call variants up to either end of the reference.

### 3.2 Active region detection

The distribution of k-mer frequencies from a set of sequence data is approximately uniform with a mean equal to the average read depth of the sample, and variants disrupt this distribution over reference k-mers. For example, a single SNP causes *k* k-mers of the sample to differ from the reference. Active region detection searches for differences between neighbors that are less likely to have occurred by chance. Then by searching for a recovery in the downstream frequencies, it attempts to resolve the left and right breakpoints in k-mer space. This region is bounded by the unaltered k-mers at its flanks, which we label *anchor k-mers.* The low-frequency and the anchor k-mers together comprise the active region.

The frequency difference between of two neighboring k-mers, *N_i_* and *N_i+1_,* is evaluated and compared to *∊,* the threshold required to trigger an active region scan (*|N_i_* − *N_i+1_| > ∊*). Because read depth can vary greatly over samples, setting this parameter to some value is unlikely to perform well on real sequence data. Therefore, Kestrel sets *∊* to some quantile, *Q_∊_,* of the |*N_i_* − *N_i+1_*| for all neighboring k-mers in the reference. The default quantile, 0.90, with an absolute minimum of 5 was found to work well in practice.

Sequencing errors, PCR duplicates, GC biases, and other factors systematically work to disrupt the uniformity of the frequency distribution. Therefore, any robust approach must be equipt with heuristics to work around such biases. Kestrel uses an exponential decay function over the recovery threshold, and it ignores short peaks in the frequency distribution.

When an active region occurs over declining read coverage, the recovery k-mer frequency may be lower than expected. As the scan moves further from the left anchor k-mer, the expected recovery frequency should be relaxed. The exponential decay function, *f*(*x*) **(Equation 2),** is employed to reduce the recovery threshold as the active region extends. *f*(0) is the anchor k-mer frequency, and it approaches a lower bound, *f_min_,* asymptotically. By default, *f_min_* = 0.55 · *f*(0) to avoid ending an active region prematurely on a large heterozygous variant region. *f*(*x*) is defined by scaling and shifting the standard exponential decay function, *h*(*x*) **(Equation 1).**

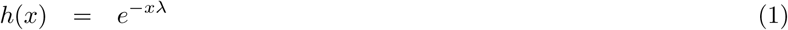

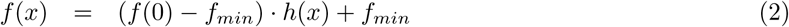

*h*(*x*) is parameterized by λ, which must be also be set, but Kestrel does not configure this parameter directly because it is difficult to know how to choose a reasonable value. Instead, λ is chosen by a configurable parameter, *α*, that is defined as the proportion of the decay range, *f*(0) − *f_min_,* after 1 k-mer **(Equation 3).** This provides a more intuitive way to define how rapidly the recovery threshold is allowed to decline.

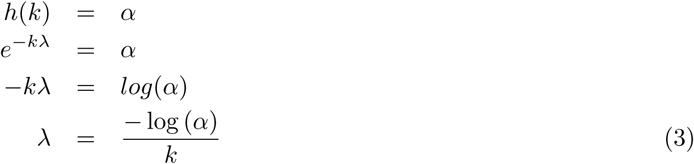

At *k* k-mers, the recovery threshold *f*(*k*) = *α* · (*f*(0) − *f_min_*) + *f_min_.* In other words, *f*(*k*) has declined in its range from *f*(0) to *f_min_* by a factor of *α.* This is true for all *nk* such that *f*(*nk*) = *α^n^* · (*f*(0) − *f_min_*) + *f_min_* **(Fig. 2).**

**Figure 2:**
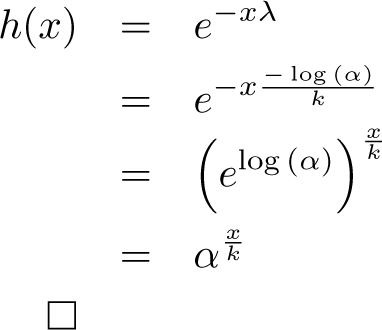
A proof of the claim that *f*(*k*) declines by 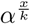. At for every *x* = *nk,* the decay function has declined *α^n^.*

### 3.3 Haplotype alignment

After the endpoints of an active region are found, the actual sequence, haplotype, from the sample must be reconstructed. This process is similar to a local assembly from k-mers, but it is implemented to keep resource usage at a minimum. Since the reference sequence over the whole region is known and anchor k-mers have been chosen, the search through k-mers in the sample can be greatly simplified.

The process begins by initializing the haplotype with the left anchor k-mer. Then by removing the left-most base of the k-mer and appending a new base to the end, the k-mer can be shifted one base to the right. Each nucleotide can be appended and the frequency checked. The base that produces a k-mer with the highest frequency is appended to the haplotype. If more than one base produces an acceptable frequency, then the alignment is split by saving the state of reconstruction with alternate bases, continuing with the highest-frequency k-mer, and returning to the alternatives after the current haplotype is built. In this way, multiple haplotypes may be built over one active region.

While this algorithm is simple enough to build the haplotype sequence, it does not know when to terminate. It could continue until the right anchor k-mer is found, but such reference-naïve reconstruction would certainly waste many CPU cycles extending erroneous sequence. Because the active region sequence and the haplotype sequences are known, an alignment could be performed as it is extended. A global alignment would be ideal except that it would have to be recomputed each time a base is added, and performance of such an algorithm would be unacceptable in all but the most trivial of applications.

An ideal algorithm would align over the active region from end to end, anchor the left ends of the haplotype and active region, and allow the right end of the haplotype to extend. To accomplish this, Kestrel employs a modified Smith-Waterman alignment on the extending haplotype. Two key modifications were made to Smith-Waterman; (i) any subalignment with a score of 0 cannot be extended, and (ii) the alignment must begin with a score greater than 0. Because of these modification, the alignment must begin with a non-zero score, and the gap extension penalty must be non-zero to prevent unbounded extension.

Smith-Waterman is a dynamic programming approach where matrices track scores using scores from shorter sub-alignments as the alignment extends (Eddy, 2004). The active region positioned over the vertical axis and the haplotype over the horizontal axis of the score matrix. We define the active region base in row *i* as *xi* and the haplotype base in column *j* as *y_j_.* As haplotype bases are added to the alignment, a column is appended to the matrix. (**Section 3.6** discusses how this is done efficiently).

Two scores (*R_match_* and *R_mismatch_*) are applied to aligned bases depending on if the bases are the same or not (respectively). For convenience, we define *match*(*i, j*) **(Equation 4)** to return *R_match_* if *x_i_* and *y_j_* match, or *R_mismatch_* if they do not. This implementation employs an affine gap model that allows for distinct gap open (*R_open_*) and gap extension (*R_gap_*) penalties. This type of model requires three matrices over *x* and *y* to track the scores. One matrix, *S_aln_,* contains scores through aligned (matched or mismatched) bases. Two more score matrices, *S_gact_* and *S_ghap_,* contain scores through gaps in the active region and gaps in the haplotype, respectively. Using these definitions, the modified Smith-Waterman score update process is defined in **Equations 5-7.**

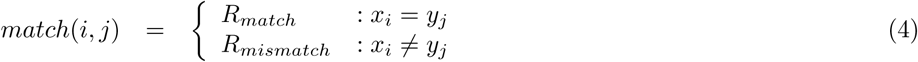

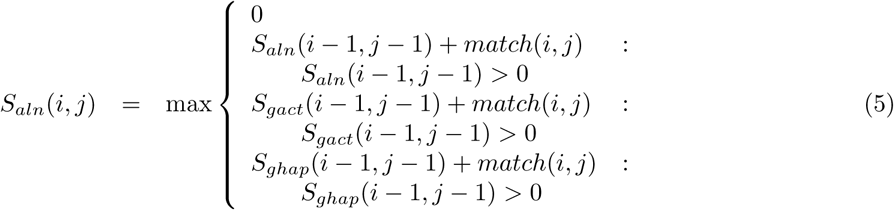

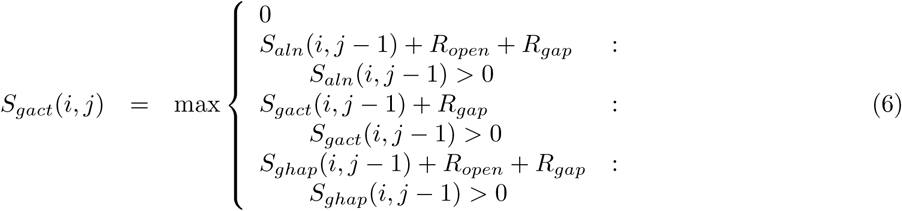

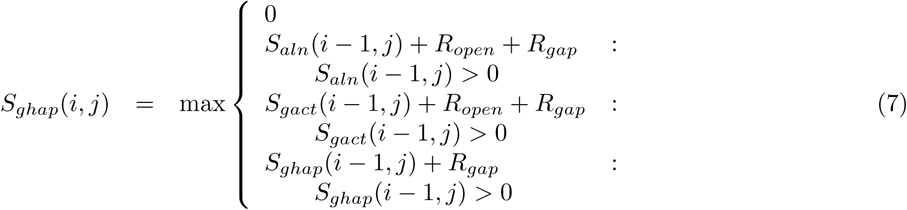

To initialize the alignment, the bases of the active region are positioned along the vertical axis of the matrices, and the each base of the anchor k-mer creates one column. The first base of the sequences is in row and column 1. Row and column 0 exist for convenience, but they do not align any bases. The initial alignment score, *R_init_* is assigned (*S_aln_*(*i, i*) = *R_init_,* 0 ≤ *i* ≤ *k*) where *k* is the size of k-mers. All other scores in all three matrices are initialized to 0. A fourth matrix, *T,* contains traceback information from the end of an alignment to *S_aln_*(0, 0). It is initialized so that *T*(*i, i*) → *T*(*i* −1, *i* − 1), 0 *< i* ≤ *k.* This initialization of *S_aln_* and *T* creates a single path for the anchor k-mer in the alignment, and all acceptable alignments must enter this path at *S_aln_*(*k*, *k*). The alignment extends from the anchor k-mer as already described.

### 3.4 Alignment termination

Since the alignment must cover all of the active region, only the last row of *S_aln_* needs to be queried to find the best score for the current alignment, *R_max_* **(Equation 8).** The maximum potential score that might be obtained by adding more bases is determined by examining the last column of *S_aln_.* The best possible score from *S_aln_*(*i*, *j*) is the case where all subsequent bases of the active region are aligned with matched based, *maxpot*(*i, j*) **(Equation 9),** and *R_maxpot_* is the maximum of *maxpot*(*i, j*) **(Equation 10).** If *R_max_ > R_maxpot_,* haplotype extension terminates.

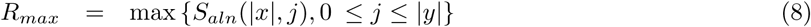

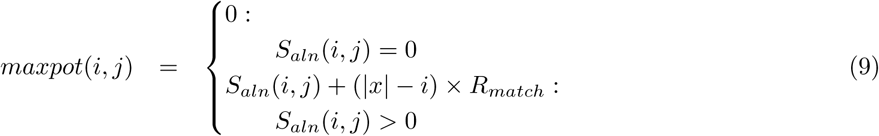

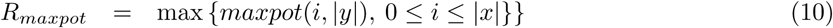

### 3.5 Variant calling

The variants are interpreted from the alignment, and the locations of the variants are translated with respect to the location of the active region. The alignment provides a convenient way to identify mismatched bases and gaps in either sequence.

If any cell of trace matrix, *T,* has more than one path out, then there is more than one optimal alignment, and so there are multiple ways to translate the alignment to variant calls. When comparing two alignments, the one with the first non-matching base is given a higher priority. If the non-matching bases agree (same variant), then the next non-matching base is queried. If the non-matching bases do not agree, then alignments are prioritized by mismatch, insertion, and deletion, in that order. This gives the algorithm predictable output for cases such as a deletion in a homopolymer repeat; Kestrel will always report that the first base was deleted even though the alignment score would be the same for a deletion at any locus of the repeat. The effect is to left-aligns variant calls.

Approximate read depth of an active region is estimated by summing the depth of all haplotypes, which may include the reference haplotype. Similarly, summing only the haplotypes that support a variant call gives the approximate depth of the variant. Because haplotypes may share k-mers, we use the minimum k-mer frequency as a conservative estimate of read depth of any single haplotype. These estimates may be useful for filtering variants with low support.

### 3.6 Alignment implementation

Because of the nature of the dynamic programming algorithm, only the last column of the score matrices (*S_aln_, S_gact_,* and *S_ghap_*) needs to be stored while the next column is built. Therefore, each of these matrices can be reduced to two arrays where one contains the last column, and one contains the new column being added. When alternate haplotypes are explored and the alignment splits, only one array for each matrix must be saved.

The traceback matrix, *T,* is more complex because it is not stored as a matrix. Instead, it is a linked-list of alignment states that always leads back to *S_aln_*(0, 0). When a non-zero score is added to a score matrix, a link is added to *T.* For each non-zero score in score matrices, a link to a node in *T* is stored.

Since *T* is a linked list that is only traversed toward *S_aln_*(0, 0), one node of the alignment may have several links into it. Therefore, different haplotypes may link to the same node in *T* where they split and no part of *T* needs to be duplicated or saved. Both haplotypes will trace back to the point where they diverged and continue toward *S_aln_*(0, 0).

The linked list structure has another more subtle property that Java uses to keep memory usage low. When an alignment path reaches a dead end, the node at the end of the path has no reference to it. This allows the Java Virtual Machine (JVM) garbage collection (GC) to detect and remove these nodes. In other words, GC can automatically prune dead branches of *T.* This improves scalabilty by reducing the memory requirements for large active regions where many haplotypes are investigated.

## 4 Results

### 4.1 *S. pneumoniae* test case

We analyzed the four penicillin binding protein (PBP) genes of *Streptococcus pneumoniae* (*S. pneumoniae)* targeted by *β*-lactam compounds. According to several studies, 20% or more of a PBP gene may be altered by inter-species recombination (Martin *et al.,* 1992; Laible *et al.,* 1991), and this can create mosaic PBP genes with a lower *β*-lactam binding affinity. Because these recombination events often alter hundreds of contiguous bases, variant calling from a standard alignment pipeline cannot characterize them.

We obtained 181 samples from 29 serotypes recently released by the Centers for Disease Control and Prevention from NCBI under BioProject PRJNA284954 (SRR2072210-SRR2072387, SRR2076738-SRR2076740). These data are whole-genome 250 bp paired-end Illumina sequence reads ranging from 15 Mbp to 1,234 Mbp (median = 323 Mbp). Selecting the best reference out of more than 10 is often necessary (Li *et al.,* 2012), however, we tested Kestrel’s ability to characterize variants using a single reference for all samples, TIGR4 (NC 003028.3).

We called all variants in four PBP genes (PBP2X, PBP1A, PBP2B, and PBP2A) using three distinct approaches. The first is a standard alignment pipeline using BWA (Li and Durbin, 2009, 2010), Picard (Institute, 2016), and GATK (McKenna *et al.,* 2010) HaplotypeCaller. The second is Kestrel. The third is a *de novo* assembly pipeline using SPAdes (Bankevich *et al.,* 2012), BWA, and SAMtools (Li *et al.,* 2009). Variants identified by the assembly where the depth of aligned contigs is 1 were defined as the true variants. HaplotypeCaller and Kestrel variant calls were compared to the true set with RTG (Cleary *et al.,* 2015) vcfeval.

Contamination in the PBP genes was identified in SRR2072298, SRR2072306, SRR2072339, SRR2072342, SRR2072351, and SRR2072379, and these samples are not included in our analysis results. For all of these samples, there were many variant calls in the PBP genes with a relative depth of 0.70 or less, which is unlikely in a monoploid organism. The size of the IKC files is also larger than expected, which supports our conclusion that there is sequence data in these FASTQ that does not belong the sample. We acknowledge but did not investigate the possibility that PBP genetic material in these samples was inserted in a plasmid. Based on the IKC files, two more samples (SRR2072345 and SRR2072352) are also likely contaminated, but we saw no evidence for this in the variant calls over the PBP genes, and these samples were included. Two other samples (SRR2072219 and SRR2072360) did not contain complete sequence data and were also removed. They contain 10 Mbp and 37 Mbp, respectively, where the median has 323 Mbp and the next lowest sample has 92 Mbp.

With the minimum k-mer frequency set to 5, sequence read errors are filtered out of the distribution of k-mers. It is then interesting to note that a sample has a finite set of representative k-mers, and once this set is reached, the IKC file ceases to grow in size. BAM files must store a record for each read, and so they grow approximately linearly with read depth. The samples suspected of contamination and low coverage indeed show an increase and a decrease, respectively, in IKC file size (**Fig. 4**).

**Figure 3:**
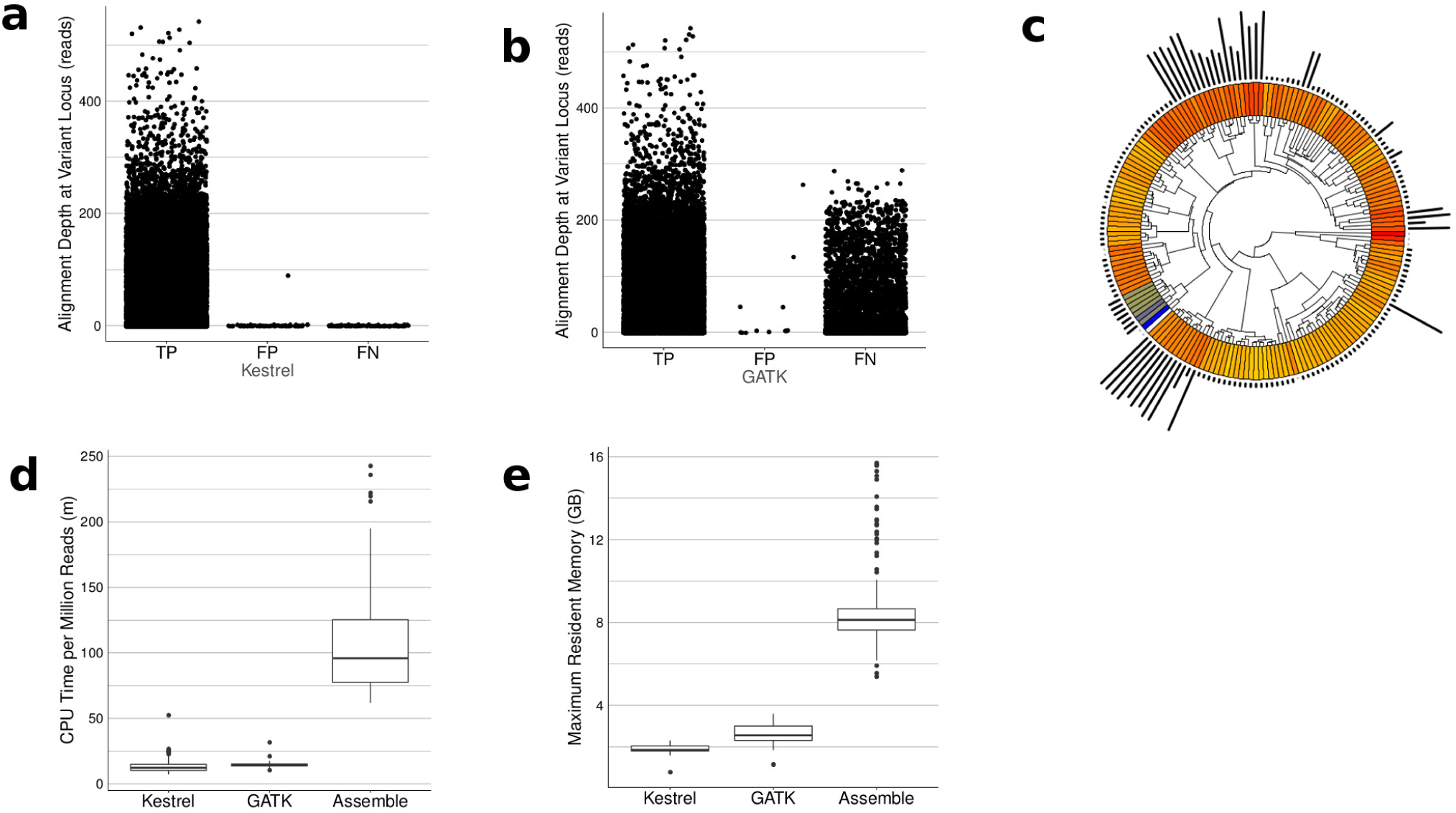
Results of testing Kestrel and the alignment approach using GATK over 173 *S. pneumoniae* samples. (**a**) Variant call summary for Kestrel calls depicting TP, FP, FN calls. (**b**) Variant call summary for GATK calls depicting TP, FP, and FN calls. (**c**) A plot depicting the phylogeny of all samples by ANI (inner track), the distance from the reference (blue) by ANI as a blue-yellow-red heatmap (middle track), and the relative number of variants in each sample (outer track). (**d**) CPU minutes per million reads. When an assembly is required, the required CPU time increases. (**e**) Maximum memory usage in gigabytes (GB) for each pipeline.

**Figure 4:**
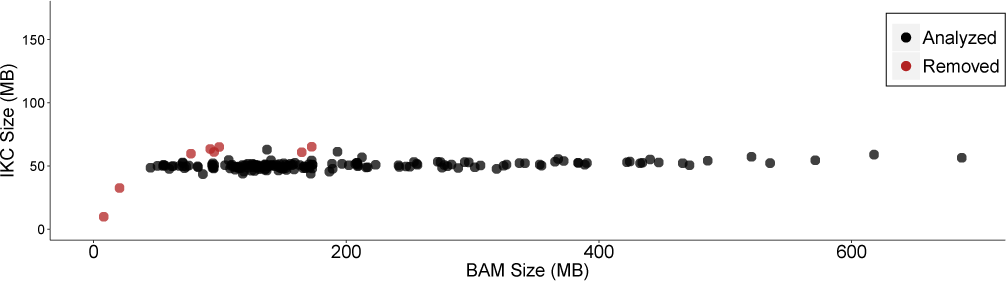
Size of BAM files vs IKC files for each sample. Since low frequency k-mers are removed, the size of an IKC file does to continue to grow with read depth once a full set of representative k-mers are present. Since BAM files have a record for each read, their size does increase with the number of reads. Samples removed for suspected contamination and low coverage are shown in red. Low-coverage samples lack a representative set of k-mers at sufficient frequency, and so the file size falls below the distribution. Samples with contamination contain k-mers that do not belong to the sample, and so their size rises above the distribution.

Kestrel produced 29,806 true positive (TP) variant calls with 73 false positive (FP) and 100 false negative (FN) calls **(Fig. 3(a))** (sensitivity = 1.00, FDR = 0.00). Because mosaic regions disrupted the alignments, GATK was only able to produce 17,636 TP variant calls with 12 FP and 11,777 FN calls **(Fig. 3(b))** (sensitivity = 0.60, FDR = 0.00). GATK tends to represent dense SNPs as insertion/deletion pairs, and so the expected true variants differs between the two approaches. These are a diverse set of samples varying in their relationship with the reference and mosaic content, and a few samples contribute most of the variants **(Fig. 3(c)).**

Kestrel required an average of 13.3 CPU minutes per million 250 bp reads (min/M-reads), the alignment approach required an average of 14.5 min/M-reads, and the assembly approach required 106.2 min/M-reads **(Fig. 3(d)).** For all steps of the pipeline, Kestrel consumed an average of 1.0 GB of memory, GATK consumed an average of 2.7 GB, and the assembly approach consumed 9.1 GB. Maximum memory consumption was 2.3, 3.6, and 15.7 GB respectively for Kestrel, GATK, and assembly **(Fig. 3(e)).** Runtime metrics were obtained on a 12 core machine (2 x Intel Xeon E5-2620) with 32 GB of RAM (DDR3-1600), RAID-6 over SATA drives (3 GB/s, 72K RPM), and CentOS 6.7.

### 4.2 MLST test case

Multilocus Sequence Typing (MLST) attempts to cluster samples by analyzing the content of some set of genes. The current method is to build a BLAST (Altschul *et al.,* 1990) database from *de novo* assembled contigs and to use the database for comparing each allele to the sequence data (Larsen *et al.,* 2012; Jolley and Maiden, 2010). A whole-genome assembly is an expensive operation, so we tested Kestrel’s ability to bypass it.

We obtained 7 *Neisseria meningitidis (N. meningitidis)* samples from ENA study PRJEB3353 (ERR193671-ERR193677) (Reuter *et al.,* 2013) and allele sequences for 7 house-keeping genes (adk, aroE, abcZ, fumC, gdh, pgm, and pdhC) from pubMLST (Jolley and Maiden, 2010). These data are whole-genome 150 bp paired-end Illumina reads. For each sample, Kestrel identified the best allele by using each allele as a reference sequence.

Sequence reads were assembled with SPAdes using default options, and contigs were used to construct a BLAST database. Each allele downloaded from pubMLST for *N. meningitidis* was used as a BLAST query, and the best allele for each gene was chosen. With the allele calls, the sequence type (ST) was identified by comparing against sequence type profiles also obtained from pubMLST. This set of alleles and the sequence type represents the expected results based on the current methods.

Sequence reads were also transformed to an IKC file with KAnalyze using 31-mers, a 15-base minimizer, a minimum k-mer frequency of 5, and a minimum allele depth of 0.50. Each allele was used as a reference sequence for variant calling. For each gene, the allele with the fewest variants was chosen as the best match. If no variants were detected for an allele, the k-mer counts over the allele reference was checked to ensure that it was present in the sequence data. The best allele matches were used to identify the ST by comparing them against types from pubMLST.

There was 100% concordance between the assembly and Kestrel methods. All samples were called with a minimum k-mer frequency of 5 except ERR193672, which was reduced to 2 to call gene pgm because of low coverage.

The assembly-based MLST approach required an average of 51.6 min/M-reads and average of 7.1 GB of memory (max 7.8 GB). The Kestrel MLST approach required 10.4 min/M-reads and 1.6 GB of memory (max 1.7 GB).

### 4.3 *E. coli* test case

We tested Kestrel’s ability to call variants on whole genomes using 309 *Escherichia coli (E. coli)* samples obtained from assembled contigs in the supplementary information published by Salipante *et al.* (2015) (Dataset S7.tar.gz). The study reports on 312 samples, but two were not found in the online dataset (upec-240 and upec-52) and one consistently caused RTG vcfeval to crash (upec-9).

The contigs were aligned to an *E. coli* K-12 reference (NC_000913.3), and variants between the reference and contig were identified. To minimize the effect of assembly errors, we used ART (Huang *et al.,* 2012) with contig sequences to simulate 150 bp paired-end reads to an average depth of 30x. Kestrel was run on the whole 4.6 Mbp reference. Variants in all regions where the contig depth was 1 were analyzed with the contig alignment variants as a set of true calls (mean 4.0 Mbp).

Variants were called using the same approach and tools as the *S. pneumoniae* contigs. No variants were called in regions where the contig alignment depth was not 1. ART (Huang *et al.,* 2012) was used to simulate the 150 bp paired-end Illumina reads to a depth of 30x over all contigs regardless of their alignment. KAnalyze and Kestrel was applied to these simulated reads with the same parameters as the *S. pneumoniae* reads (k-mer size 31, quality 30, min depth 5) except the active region size, which was set to 7x the maximum size of an insertion variant, as calculated by the alignment score parameters. An additional parameter was given to Kestrel to discard variants over ambiguous reference bases, such as N, so that RTG vcfeval had an equivalent set of variants from both approaches. Unlike the *S. pneumoniae* experiment, variant calling was done over the whole genome reference instead of targeted regions. With both variant call sets, RTG vcfeval was used to test the calls.

In addition to a VCF, Kestrel can output a SAM file of the haplotypes it assembles and aligns. The SAM feature was enabled for this experiment so that the locations of haplotypes could be identified. Variant statistics were calculated twice; once over all regions where the alignment depth was one and once for all regions where there was an assembled haplotype with at least one variant. The current implementation of Kestrel discards active regions with one wildtype haplotype, so the second analysis only applies to regions where at least one variant was identified.

Over the whole genome, Kestrel yielded and FDR of 0.01, but a sensitivity of only 0.71. Kestrel must limit the size of active regions or the haplotype alignments will consume too much memory. If we examine only regions where Kestrel had an active region (mean 2.3 Mbp), the sensitivity rose to 0.98, and the FDR remained 0.01.

## 5 Discussion

The Kestrel algorithm is a framework for calling variants from sequence data using evidence found only in k-mer space. It provides a mechanism for detecting regions of variation, a method for resolving the variation to variant calls, and a set of heuristics that make it work with real data. Inference on k-mers has been applied to RNAseq analysis (Patro *et al.,* 2014; Bray *et al.,* 2016), metagenomics (Wood and Salzberg, 2014), phylogenetics (Gardner and Slezak, 2010; Gardner and Hall, 2013), and many other problems in bioinformatics. To our knowledge, Kestrel is a first-in-class variant calling implementation using only k-mers.

Although such an approach is unorthodox with respect to current standards, it is able to capture variation in regions where alignments are too noisy. In these cases, a k-mer approach can displace alternatives approaches that require far more computing resources and time to complete, such as *de novo* assemblies. Extending this approach to other domains where assemblies are the current standard, such as MLST, can also have a dramatic effect computing resources.

Despite several advantages, a purely k-mer-based approach such as this does have limitations. The most significant of these is that paired-end and sequence read context is lost when shredding sequence reads into k-mers. This information is very valuable for resolving variants in repetitive, duplicated, or low complexity regions of the genome. Some evidence of these events is still present in k-mer space, such as frequency peaks, and although Kestrel attempts to work through these events, alignments or assemblies will often have higher accuracy.

As a practical tool, Kestrel is a fast alternative to existing methods when applied to specific regions of the genome. We expect it to find use as a targeted tool or an orthogonal approach for other callers. In its current form, we cannot purport it to be a whole-genome alternative for software like Cortex or Platypus. Future developments may improve the limitations of this approach, or the algorithm may find use in a hybrid tool making use of k-mers and other data. For example, the GATK HaplotypeCaller might fall-back to such a k-mer method to rescue variants in a region where alignments suffer because of dense SNVs or large indels.

In many cases, algorithms that are free of alignments and assemblies can reduce the demand on computing resources. As high-volume sequencing technology becomes faster and cheaper, reducing the cost of data storage and analysis becomes critical (Köser *et al.,* 2012; Sboner *et al.,* 2011). Kestrel’s contribution was only marginal for calling variants where alignments would suffice, but its ability to perform analysis without more costly methods gives it a significant advantage. The computing resources Kestrel requires is well within the capabilities of a modern laptop, which is far less expensive than the powerful machines or computing clusters that would be required to do the same work in the same amount of time. Such a cost savings may conserve vital research funds or enable routine analysis where funding and equipment are not as readily available.

## 7 Acknowledgements

Ben Metcalf and Bernard Beall from the Centers for Disease Control and Prevention provided data and insight for the *S. pneumoniae* experiment. Dawn Audano designed the graphic for **Fig. 1.**

